# Delaying Cancer Progression by Integrating Toxicity Constraints in a Model of Adaptive Therapy

**DOI:** 10.1101/2025.04.24.650205

**Authors:** Jana L. Gevertz, Harsh Vardhan Jain, Irina Kareva, Kathleen P. Wilkie, Joel Brown, Yitong Pepper Huang, Eduardo Sontag, Vladimir Vinogradov, Mark Davies

## Abstract

Cancer therapies often fail when intolerable toxicity or drug-resistant cancer cells undermine otherwise effective treatment strategies. Over the past decade, adaptive therapy has emerged as a promising approach to postpone emergence of resistance by altering dose timing based on tumor burden thresholds. Despite encouraging results, these protocols often overlook the crucial role of toxicity-induced treatment breaks, which may permit tumor regrowth. Herein, we explore the following question: would incorporating toxicity feedback improve or hinder the efficacy of adaptive therapy? To address this question, we propose a mathematical framework for incorporating toxic feedback into treatment design. We find that the degree of competition between sensitive and resistant populations, along with the growth rate of resistant cells, critically modulates the impact of toxicity feedback on time to progression. Further, our model identifies circumstances where strategic treatment breaks, which may be based on either tumor size or toxicity, can mitigate overtreatment and extend time to progression, both at the baseline parameterization and across a heterogeneous virtual population. Taken together, these findings highlight the importance of integrating toxicity considerations into the design of adaptive therapy.

## 1 Introduction

Systemic anti-cancer therapies, including chemotherapy, targeted therapies, immunotherapies, and hormone treatments, are essential components of modern oncology. In patients with metastatic disease, where cure is often elusive, the primary goals are to control tumor growth, extend survival, minimize treatment-related toxicities, and alleviate cancer-associated symptoms (1). Typically, a maximum tolerated dose (MTD) protocol is used to achieve these goals. The goal of MTD is to eliminate as many cancer cells as possible by administering the drug at the highest dose deemed safe until toxicity limitations are reached.

Despite significant advances in the precision and effectiveness of systemic anti-cancer therapies, two persistent challenges remain. The first is the management of treatment-induced toxicity (2–4). Toxicities associated with MTD treatment include fatigue, gastrointestinal side effects, and hematological complications (5–9). These toxicities adversely affect patient quality of life and often necessitate dose reductions, treatment interruptions, or even discontinuation, all of which ultimately undermine therapeutic effectiveness.

The second persistent challenge of MTD therapy is the emergence of therapeutic resistance. In clinical practice, a tumor is considered sensitive if it shrinks or remains stable with treatment, and resistant if it continues to grow. However, because tumors are inherently heterogeneous, they typically contain subpopulations of cells with varying treatment sensitivities (10). Under the sustained treatment typical of MTD therapy, the elimination of sensitive cells can create an environment in which resistant clones proliferate freely. This phenomenon, known as competitive release, accelerates resistance-driven progression and diminishes treatment efficacy over time (11,12).

Adaptive therapy has emerged as a promising alternative to traditional MTD approaches. Rather than aiming for complete tumor eradication, adaptive therapy strategically modulates dosing schedules to preserve a population of sensitive cells that can suppress resistant clones through intra-tumor competition (13,14). Its implementation is governed by clinical decision rules based on measurable indicators such as tumor size or circulating biomarkers. For example, a dose-skipping regimen administers a high dose until a predetermined tumor response (e.g., a 50% reduction in size) is achieved, and then pauses treatment until the tumor regrows to a defined threshold, often its initial size. Alternatively, a dose-modulation strategy adjusts treatment doses incrementally – increasing or decreasing them on the basis of the tumor’s response over time (15).

The effectiveness of adaptive therapy can be enhanced if resistance comes with a fitness cost. This occurs when the mechanisms conferring resistance, such as the upregulation of energy-intensive drug efflux pumps, reduce a cell’s overall fitness in the absence of therapy (16–18). Indeed, adaptive therapies have already been tested experimentally (19–21) and are currently being applied across multiple clinical trials (<u>NCT02415621</u>; <u>NCT03511196</u>; <u>NCT03543969</u>; <u>NCT03630120</u>).

However, a crucial, yet underexplored, aspect of adaptive therapy is the impact of treatment-induced toxicity on dosing decisions. Toxicity constraints can limit treatment frequency or intensity, potentially allowing both sensitive and resistant cell populations to proliferate during treatment interruptions. This could disrupt the competitive dynamics that are essential to adaptive therapy’s success. Equally, integrating toxicity feedback into dosing strategies could improve patient tolerability and enable prolonged treatment, ultimately enhancing outcomes.

Attempts to model toxicity which can be found in the literature mainly focus on white blood cell counts and neutropenia induced by chemotherapy (22–26). In (27), the authors model chemo-induced toxicity effects as a loss of muscle mass, and in (28), the authors model toxicity effects via both weight loss and a comprehensive index that takes into account 57 side effects. Toxicity has also been included as an additional constraint in the design of optimal control dosing strategies (29,30).

Approaches to model toxicity include an assumption that the effect is present and constant during the whole treatment phase (22,23), that it is proportional to drug concentration (25,29,30), or that it is proportional to drug and tumor burden (28). In (24), the toxic effect of the drug directly reduces the neutrophil proliferation rate via an inverse polynomial form of the drug concentration. In (22), a stochastic model is introduced wherein toxicity is modeled as an increase in the death rate of white blood cells. Through a virtual clinical trial, the authors demonstrate that managing toxicity throughout treatment by modulating or pausing doses does not compromise overall treatment outcomes. In (31), the authors use a modified logistic competition model with sensitive and resistant cells to show that in numerical simulations continuously-dosed adaptive therapy outperforms discretely-dosed adaptive therapy, in terms of both time to disease progression and controlling toxicity. Their results assume toxicity is a nonlinear function of drug dose satisfying specific conditions (such as being strictly increasing and concave up).

Previously considered control problems (30) introduce toxicity as a constraint, but do not monitor the emergence of drug resistance or the strategy of adaptive therapy. Dose interruptions in a model of resistance is presented in (22), but in that paper, toxicity is considered as a constant effect that reduces white blood cell growth, and is not dependent on drug concentration or accumulated exposure. In (31), where the authors consider continuous or intermittent adaptive therapies to manage emergence of resistance, toxicity is modelled as a side effect that can be used to determine when too much drug has been administered but is not included in dosing decision making.

In this work, we explore the following critical question: would incorporating toxicity feedback improve or hinder the efficacy of adaptive therapy? To address this question, we propose a novel mathematical framework for including toxicity both in the decision-making process of daily therapy, and in the formulation of adaptive regimes based on maintaining the tumor size within a pre-specified range. Our simulations, designed to mimic a theoretical murine study with an unspecified chemotherapeutic agent, compare the impact of toxicity constraints on time to progression (TTP) for daily treatment versus adaptive therapy schedules. Our results demonstrate that the degree of competition between sensitive and resistant populations, along with the growth rate of resistant cells, critically modulates how TTP is impacted by the incorporation of toxicity feedback. Notably, our model identifies circumstances where strategic treatment breaks, which may be based on either tumor size or toxicity, can mitigate overtreatment and extend TTP, both at the baseline parameterization and across a virtual population. We conclude with a discussion of the model’s limitations and propose future directions aimed at optimizing cancer treatment strategies to simultaneously address resistance and toxicity, thereby improving patient quality of life.

## 2 Model Development

To investigate the relationship between adaptive therapy and toxicity, we introduce a novel mathematical model that integrates drug pharmacokinetics and toxicity dynamics into an existing framework of competitive interactions between treatment-sensitive (*S*(*t*)) and treatment-resistant (*R*(*t*)) cancer cells, compare to (Strobl et al. 2021). In this model, it is assumed that the two subpopulations compete for, and grow logistically up to, a shared carrying capacity *K*. Sensitive cells can be killed by the drug, while resistant cells cannot. Drug concentration *C*(*t*) is described by a simple one compartment pharmacokinetic model with linear clearance. Toxicity *T*(*t*) increases with drug concentration *C*(*t*), and resolves at a fixed linear rate.

The resulting system of equations is as follows:

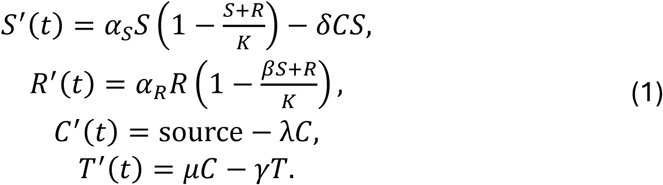

Here *α*_*S*_ and *α*_*R*_ are the growth rates of the sensitive and resistant cells, respectively, and *β* is the competition suppression factor that sensitive cells impose on resistant cells. The drug is assumed to be instantaneously injected into the model through the source term, has a clearance rate of *λ*, and a killing efficacy of *δ* on the sensitive cells. Toxicity accumulates proportionally to the drug concentration with parameter *μ*, and decays exponentially with parameter *γ*. Baseline values of parameters, together with their meanings, are summarized in Table 1. We remark that in our formulation, resistance to drug incurs a cost so that resistant cells cannot proliferate faster than sensitive cells. Mathematically, we express the growth rate of resistant cells as: *α*_*R*_ = *ϵα*_*S*_, where 0 ≤ *ϵ* ≤ 1.

**Table 1.** Baseline parameter values used in model system. (**1**).

| Parameter | Description | Value | Dimensions |
| --- | --- | --- | --- |
| $\alpha_S$ | Growth rate of sensitive cancer cells | 1 | per day |
| $\alpha_R$ | Growth rate of resistant cancer cells | $\epsilon \alpha_S$ | per day |
| $\epsilon$ | Dimensionless constant between 0 and 1 | 0.4 | dimensionless |
| $K$ | Shared carrying capacity | 100 | dimensionless |
| $\delta$ | Rate of drug-induced cancer cell death | 1 | per day |
| $\beta$ | Competition parameter between sensitive and resistant cancer | 2.4 | dimensionless |
| $\lambda$ | Drug clearance rate | 0.693 | per day |
| $\mu$ | Rate of toxicity accumulation | 1 | per day |
| $\gamma$ | Toxicity recovery rate | 0.4 | per day |
| $Rx_{\text{off}}$ | Efficacy threshold triggers therapy pause if total tumor size falls below this value | 40% of initial tumor size | dimensionless |
| $Rx_{\text{on}}$ | Efficacy threshold triggers therapy restart if total tumor size surpasses this value | Initial tumor size | dimensionless |
| $Tox_{\text{off}}$ | Toxicity threshold triggers therapy pause if toxicity surpasses this value | 2 | dimensionless |
| $Tox_{\text{on}}$ | Toxicity threshold triggers therapy restart if toxicity falls below this value | 1 | dimensionless |
| **Treatment Failure** | Tumor size defining treatment failure and disease progression | 75 | dimensionless |

In the interest of generality and wide applicability—and because we are not modeling any specific cancer type, drug, or experimental dataset—we rescale our model variables as follows. The tumor cell populations *S*(*t*) and *R*(*t*) are expressed as fractions of an assumed carrying capacity of 100, making them interpretable as percentages of sensitive and resistant tumor cells. The drug concentration *C*(*t*) is dimensionless, reflecting its role as a generic chemotherapeutic agent. Likewise, toxicity *T*(*t*) is measured in arbitrary units to encompass a broad range of potential side effects. These rescalings provide a flexible framework within which we can explore treatment dynamics across various scenarios. The baseline parameters were selected to demonstrate the dynamical differences between the four protocols detailed in Section 2.2 (Defining Treatment Protocols).

Model (1) is solved in MATLAB® using ode23, a stiff solver that implements a modified Rosenbrock formula of order two. Unless otherwise indicated, the scripts used to solve and analyze the toxicity model are available at https://github.com/jgevertz/toxicity.

### 2.1 Defining Treatment Failure

Following the RECIST (Response Evaluation Criteria in Solid Tumors) criteria used in clinical practice – which typically defines disease progression as at least a 20% increase in tumor size from some baseline (32) – we define therapy failure as the total tumor burden rising 50% above its value at treatment initiation. In our model, the tumor burden starts at 50% of the carrying capacity, and treatment failure is thus defined as a tumor burden of 75% of the carrying capacity (that is, the initial tumor burden is 50, and treatment failure occurs when *S* + *R* ≥ 75). TTP is defined as the time from treatment initiation until the tumor burden reaches the treatment failure size. We emphasize that in a clinical setting, an oncologist would not know the tumor’s carrying capacity and, instead, would make decisions based on measurable tumor burden obtained via imaging or other methodologies. To reflect this, efficacy-based decisions in our model are related to the tumor size at therapy initiation rather than to its (unknown) carrying capacity. In our numerical simulations, the tumor’s initial composition is set at 99% sensitive cells and 1% resistant cells at 50% carrying capacity, or *S*(0) = 49.5 and *R*(0) = 0.5. Treatment starts on day 1, so both the drug concentration and toxicity are assumed to be zero initially, that is, *C*(0) = *T*(0) = 0.

### 2.2 Defining Treatment Protocols

Next, we present our approach to integrating both efficacy-based (tumor size) and toxicity-based (drug-induced toxicity level) feedback into the decision rules for starting, pausing, and resuming treatment. The resulting four protocols are summarized in Figure 1.

**Figure 1.**
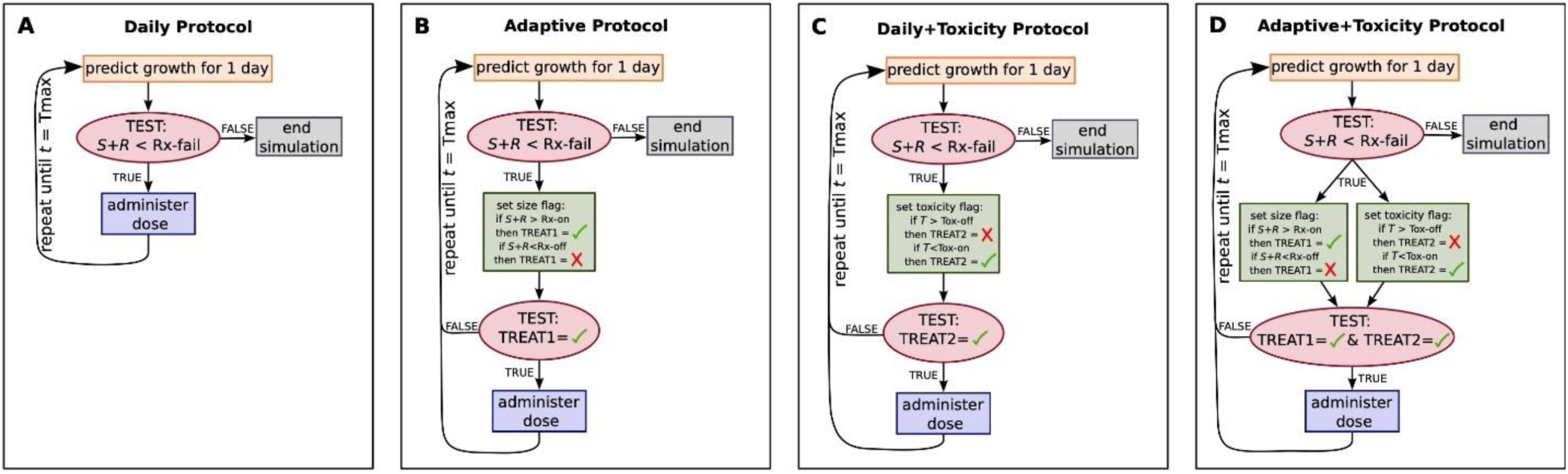
Summary of the four proposed protocols: (A) daily protocol, (B) adaptive protocol with tumor size-driven treatment cycles, (C) daily protocol with toxicity-driven treatment cycles, and (D) adaptive protocol with both tumor size- and toxicity-driven treatment cycles.

#### Daily Protocol

A fixed dose of 1 unit is administered at daily intervals (time *t* = 1 day) until either treatment failure or the total elapsed time hits 100 days, whichever occurs first. Tumor burden is monitored daily, just before treatment administration, to check for disease progression. This protocol is summarized in Figure 1A.

#### Adaptive Protocol

Treatment is administered in daily 1 unit doses until the total tumor size, *S*(*t*) + *R*(*t*), falls below the Rx_off_ threshold. At this point, therapy is paused until the tumor size regrows and again exceeds the upper Rx_on_ threshold, prompting treatment to resume. Similar to the daily protocol, tumor burden is monitored daily, just before potential treatment administration, to determine if disease progression has occurred. This protocol is summarized in Figure 1B.

#### Toxicity Feedback

This protocol can be incorporated into either of the previous two protocols described above, thus acting as an override switch for pausing therapy. Specifically, if toxicity levels are above a designated threshold Tox_off_ when the next treatment decision is to be made, then treatment is halted regardless of the tumor size. Therapy can only resume when the toxicity level at the time of the next treatment decision drops below the Tox_on_ threshold. In the daily protocol with toxicity feedback, dosing (re)starts when toxicity is below Tox_on_, and will continue until toxicity rises above Tox_off_. In the adaptive protocol with toxicity feedback, however, treatment (re)starts when both toxicity is below Tox_on_ and the tumor size is above Rx_on_. Treatment will pause when either toxicity is above Tox_off_ or the total tumor size is below Rx_off_. These conditions are tested daily, prior to dose administration. The cycles will continue until treatment failure or total simulation time hits 100 days. Figures 1C and 1D summarize adding toxicity feedback to the daily and adaptive protocols, respectively.

Each administered dose is assumed to be 1 unit of drug, and the potential treatment period is simulated over 100 days. During this time, the simulation may or may not predict disease progression by day 100. Time to progression (TTP) and tumor composition at TTP are recorded at the end of each simulation. If the tumor did not progress in the 100-day simulation, then TTP is recorded to be 150 (a randomly selected value greater than 100 to indicate treatment did not fail during the simulation time). Simulations will follow one of the four treatment protocols described above.

### 2.3 Global Sensitivity Analysis Framework

To assess the robustness of model predictions to variability in parameter values, we conduct a global sensitivity analysis using the extended Fourier Amplitude Sensitivity Test (eFAST). eFAST is a variance-based decomposition technique capable of efficiently handling nonlinear models (33,34). The method quantifies the sensitivity of a model’s output to variations in input parameters, by computing both a first-order and a total-order sensitivity index. The sensitivity of a particular input, the ratio of the total variability in the output attributed to changes in that input, is found by averaging over all other inputs. Parameters with high sensitivity indices are identified as influential to the model output, while those with low sensitivity indices may be regarded as negligible. The template used for eFAST implementation can be found here: http://malthus.micro.med.umich.edu/lab/usadata/

Herein, we assess the global sensitivity of TTP across the four treatment protocols for the following model parameters over the following ranges: the sensitive cell growth rate (0.5 ≤ *α*_*S*_ ≤ 1.5), the resistant cell growth rate relative to the sensitive cell growth rate (0.2 ≤ *ϵ* ≤ 1, *α*_*R*_ = *ϵα*_*S*_), the rate of drug-induced cancer cell death (0.5 ≤ *δ* ≤ 1.5), the competition parameter (0.5 ≤ *β* ≤ 3.5), and the toxicity recovery rate (0.2 ≤ *γ* ≤ 0.8). The output of this sensitivity analysis then informs a set of parameter sweeps, wherein we quantify the impact that simultaneously varying two model parameters have on TTP in each of the four treatment protocols.

### 2.4 Defining Virtual Patient Framework

To extend our numerical simulations and two-dimensional parameter sweeps to better understand variability across parameter space, we utilize a virtual population approach (35). In our computational framework, each virtual patient (VP) in the virtual population is represented by a set of parameter values related to characteristics of tumor growth and response to treatment. That is:

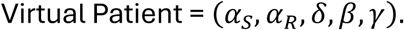

The baseline parameter values listed in Table 1 can be thought of as corresponding to an ‘average’ patient. To construct a VP, a value for each parameter in the set (*α*_*S*_, *α*_*R*_, *δ*, *β*, *γ*) is drawn as a simple random sample from a corresponding lognormal distribution. The distribution’s shape parameters *μ* and *σ*, are chosen such that the peak approximately occurs at the nominal VP parameter value reported in Table 1, and the width matches the range used in the global sensitivity analysis. This process is repeated 100 times to form a heterogeneous virtual population, or 100 sets of the 5 VP parameters.

The carrying capacity *K*, rate of drug clearance *λ*, and rate of toxicity build up *μ* are fixed across all VPs. This decision was made to preserve the consistency of key modeling assumptions and to prevent confounding of sensitivity results due to parameter interdependencies. Since tumor size at treatment initiation was set at 0.5*K* and treatment failure at 0.75*K*, varying *K* would not affect time to progression. We fixed *μ* because of its relation with *γ* – higher toxicity build-up rates *μ* can be offset by correspondingly higher toxicity clearance rates *γ*. Likewise, the drug’s cell-killing effect can be preserved even with a high clearance rate *λ* if the rate of treatment-induced cell death *δ* is also high.

## 3 Results

In this section, we evaluate the impact of four proposed protocols (see Figure 1) on treatment dynamics and time-to-progression (TTP): (i) daily protocol, (ii) adaptive protocol, (iii) daily protocol with toxicity feedback, and (iv) adaptive protocol with toxicity feedback. In all simulations, parameters are fixed at their baseline values from Table 1, except for those being explicitly varied in a parameter sweep, with the same normalized dose administered across all treatment protocols. We monitor tumor composition (fractions of sensitive and resistant cells), drug concentration, and accumulated toxicity. Protocol effectiveness is assessed based on TTP – the longer a tumor takes to fail treatment, the more effective the protocol is deemed to be.

### 3.1 Addition of Toxicity Constraints Can Extend TTP

Figure 2 displays representative model dynamics from a simulated mouse experiment for comparing the four treatment protocols. Each row corresponds to a protocol, with tumor cell time-courses plotted in the left panels, and drug concentration and toxicity time-courses in the right panels. Periods of drug administration are shaded in gray, and simulations run until the disease progresses. For each protocol, representative examples of model dynamics at alternative parameterizations are shown in Figure S1.

**Figure 2.**
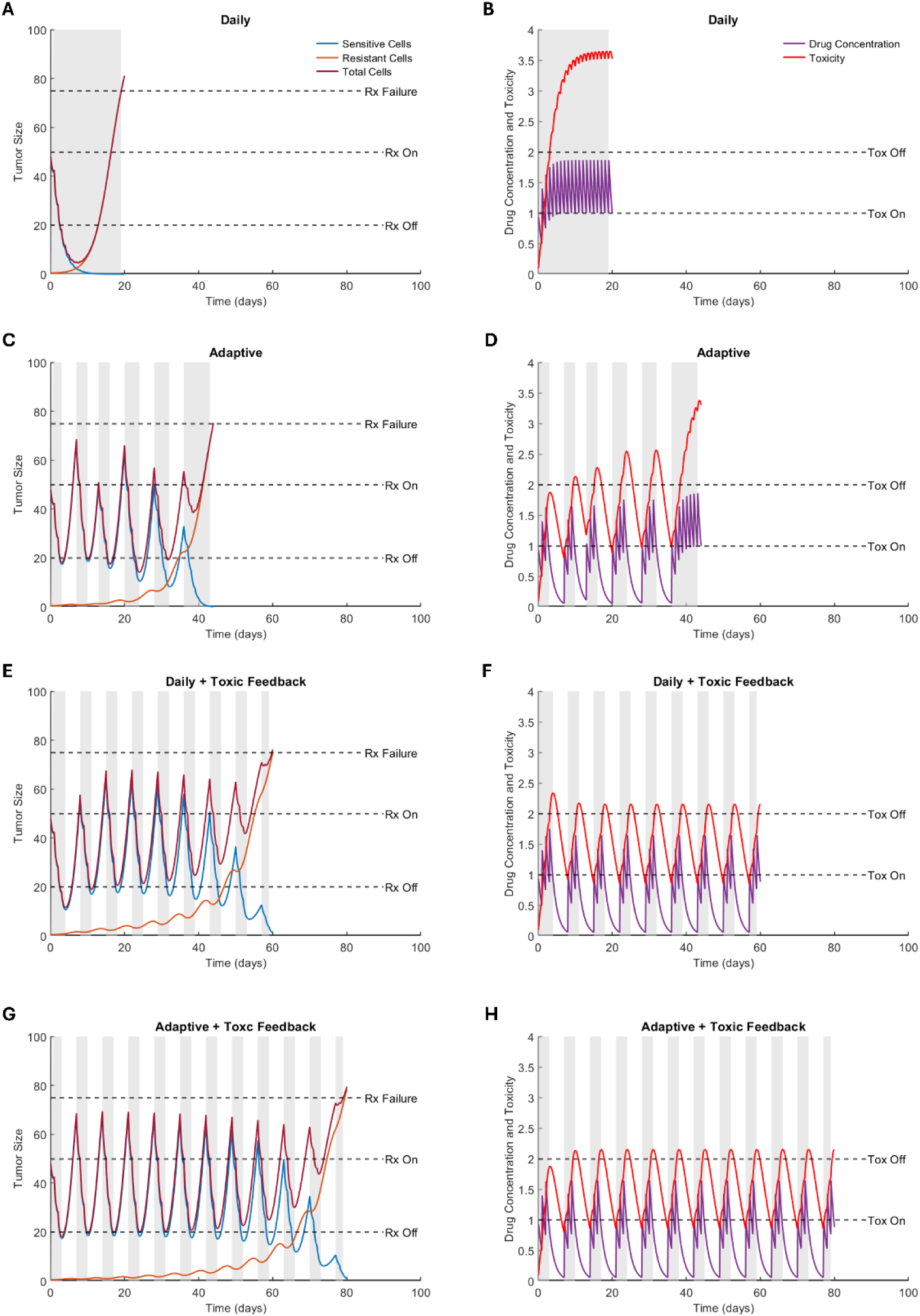
The four treatment protocols and resulting model outputs. The left column shows the time course of the total tumor size, along with the sensitive and resistant subpopulations. The right column shows the dynamics of drug concentration and accumulated toxicity. Compared protocols are daily (A-B), adaptive (C-D), daily with toxicity feedback (E-F), and adaptive with toxicity feedback (G-H). Time periods shaded grey represent when the treatment is on.

In the daily protocol (Figure 2, top row), the drug rapidly depletes sensitive cells (blue curve), resulting in an initial reduction in total tumor burden. However, this depletion removes intra-tumor competition, allowing resistant cells (red curve) to proliferate unchecked, leading to swift treatment failure. High and sustained drug exposure results in persistently elevated toxicity throughout treatment. This protocol yields the shortest TTP (19.2 days), creating a “lose-lose” scenario where resistance emerges quickly, and toxicity remains critically high.

In the adaptive protocol, tumor size-based feedback modulates treatment cycles, alternating between drug administration and off-treatment periods (Figure 2, second row). This strategy allows sensitive cells to recover during treatment pauses, enabling them to compete with and temporarily suppress resistant cells, thereby delaying treatment failure. Periodic drug holidays also reduce toxicity compared to the daily protocol, though overall toxicity continues to increase with time. Adaptive therapy extends TTP to 44 days, which is more than doubling the duration achieved under the daily protocol. However, as resistant cells gradually escape suppression by the sensitive cell population, the adaptive protocol ultimately loses effectiveness.

Incorporating toxicity-based dose adjustments into the daily protocol introduces periodic drug holidays (Figure 2, third row). These treatment interruptions, designed to allow recovery from drug toxicity, also enable partial regrowth of sensitive cells. Interestingly, for our baseline parameter values, this approach achieves a longer TTP (59.8 days) than even the adaptive protocol. This is because the toxicity feedback effectively mimics an adaptive strategy with higher Rx_on_ thresholds; that is, treatment resumes only after the tumor has grown beyond a threshold that would have triggered re-administration in the adaptive protocol.

The adaptive protocol with toxicity modulation integrates both tumor size-based and toxicity-based treatment decisions (Figure 2, bottom row). This doubly adaptive approach results in sustained on/off treatment cycles. Like the daily protocol with toxicity feedback, tumor sizes overshoot the Rx_on_ threshold because treatment cannot resume until toxicity has declined sufficiently. This has the dual effect of reducing drug exposure – and consequently, toxicity – whilst preserving the sensitive cell population during early cycles. As a result, this protocol achieves the longest TTP (79.1 days).

### 3.2 Sensitivity Analysis

Next, we perform a global sensitivity analysis using eFAST to quantify the impact of parameter variations on time-to-progression (TTP) across the four treatment protocols. The results are summarized in Figure 3. It should be noted that, eFAST assigns a minimal sensitivity value to all parameters, including dummy parameters, making the *p*-values (indicated by asterisks in each plot) crucial for interpreting the significance of each sensitivity index.

**Figure 3.**
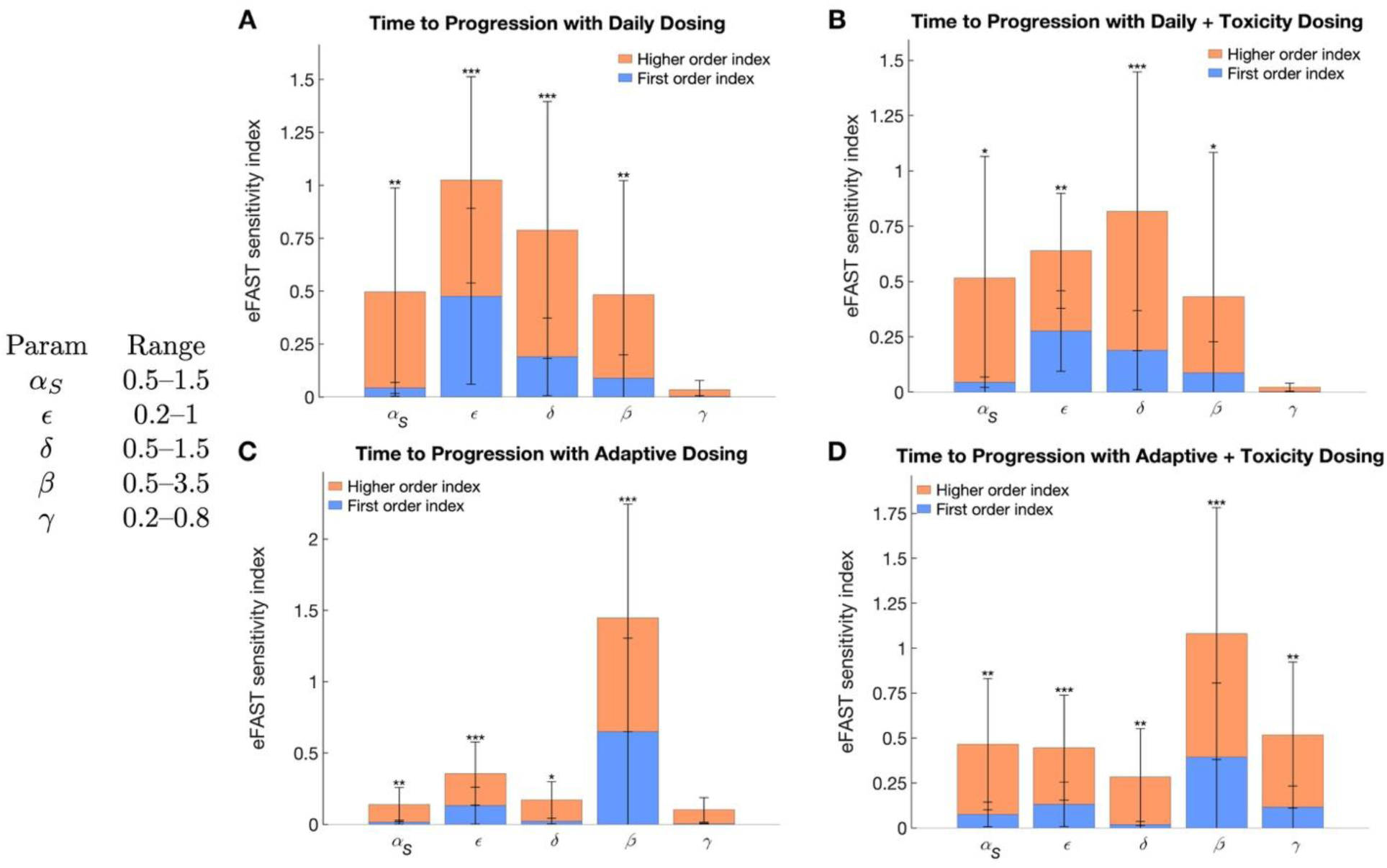
eFAST sensitivity indices as quantified by time to progression (TTP) for (A) daily protocol, (B) daily protocol with toxicity, (C) adaptive protocol, and (D) adaptive protocol with toxicity. Asterisks indicate statistical significance: * p < 0.05, ** p < 0.01, *** p < 0.001. Parameter ranges used are indicated in the graph. Note that *α*_*R*_ = *ϵα*_*S*_.

The sensitivity indices plotted in Figure 3 indicate that no single parameter strongly influences TTP in all four protocols. In the adaptive protocols (Figure 3C,D), the competition parameter β most significantly influences TTP, as indicated by having the largest first order (blue bars), and higher order (orange bars), sensitivity indices. In the daily protocols (Figure 3A,B), the parameter with the most influence TTP is the relative growth rate of resistant cells relative to sensitive cells (*ϵ*, *α*_*R*_ = *ϵα*_*S*_). Counterintuitively, the rate of toxicity recovery, *γ*, is either the least or second least sensitive parameter in any of the protocols. We explore this further in Section 3.3.

### 3.3 Effect of Parameters on TTP

Parameters *β* and *ϵ* (or *α*_*R*_ as *α*_*R*_ = *ϵα*_*S*_) were identified as significant for most protocols in Figure 3, and so we start our analysis with them. We first investigate the effect on TTP of varying *β* and *α*_*R*_ over the ranges defined in the global sensitivity analysis, with all other parameters fixed at their baseline values from Table 1. For each pair of values (*β*, *α*_*R*_) we compute the TTP for each protocol (Figure S2). The results of these four parameter sweeps are summarized in Figure 4. In particular, Figure 4A indicates which protocol(s) result in the longest TTP, and Figure 4B indicates the actual time to progression for the protocol(s) that maximized TTP. Recall that the simulation is run for 100 days. We assign a value of 150 to any simulation that does not reach treatment failure within this simulation time.

**Figure 4.**
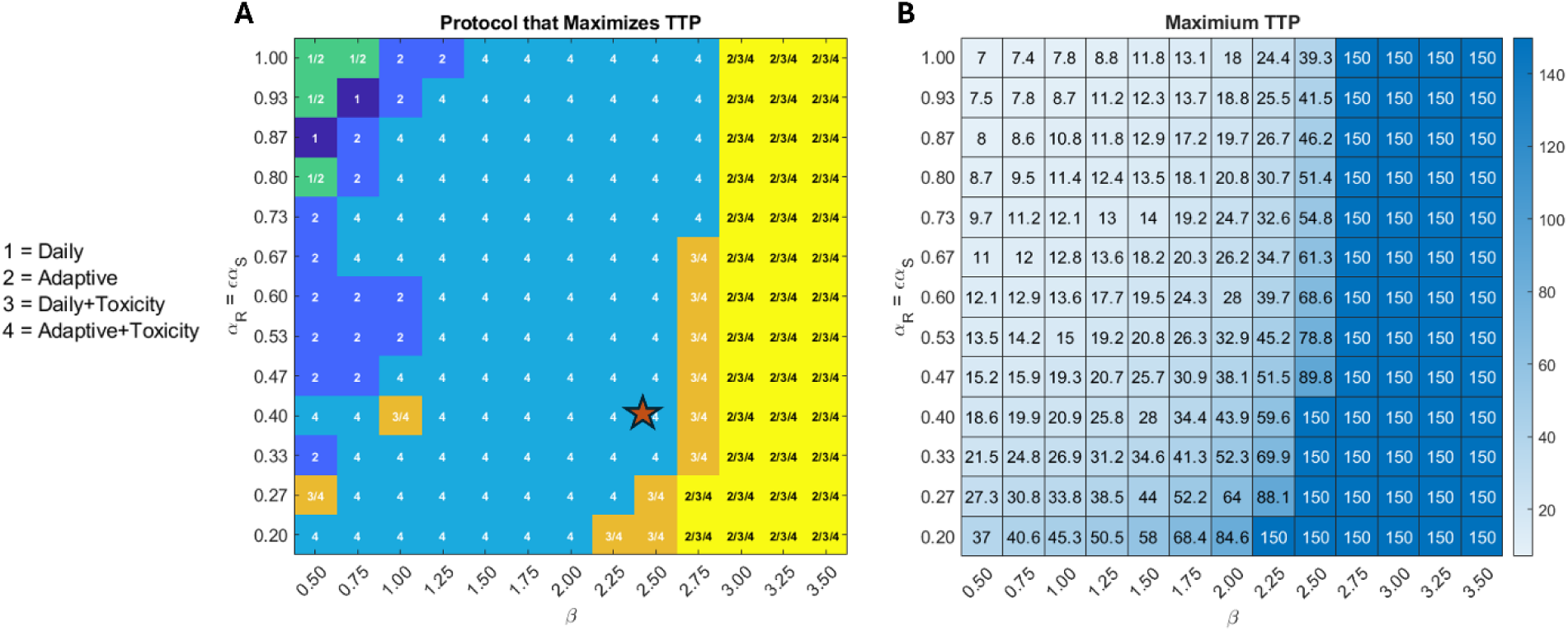
Parameter sweeps over *β* − *α*_*R*_ space. (A) Protocol that maximizes time to progression for the corresponding pair of parameter values. (B) Maximum time to progression for the optimal protocol(s) identified in (A). A value of 150 indicates that the predicted tumor did not progress within the simulation time of 100 days and thus, that the treatment protocol did not (yet) fail. The baseline parameterization is indicated with a red star. Parameters that are not varied are fixed at their baseline value in Table 1.

As shown in Figure 4A, one of the two adaptive protocols is overwhelmingly likely to maximize TTP when the resistant cells have a competitive advantage over the sensitive cells (*β* < 1). The only exception to this is when resistant cells grow at nearly the same rate as the sensitive cells (*α*_*R*_ near 1). In this scenario, the competitive advantage of the resistant cells is so extreme that none of the protocols can contain the tumor, and any stopping of treatment decreases TTP. Figure 4B shows that as the competitive advantage of the resistant cells increases (smaller *β*, larger *α*_*R*_), the optimal protocol becomes less effective. As resistant cells lose their direct competitive advantage over sensitive cells (in particular for 1 ≤ *β* ≤ 2), the incorporation of toxicity feedback into the adaptive protocol yields the optimal result, though treatment still fails within the simulation period. Finally, when sensitive cells have a very strong competitive advantage over resistant cells (*β* ≥ 2.75) or a sufficiently strong competitive advantage (2.25 ≤ *β* ≤ 2.5) coupled with significantly weaker growth of resistant cells relative to sensitive cells (small *ϵ*), TTP extends beyond the simulation window. As we move towards the parameter regime where the tumor does not progress (higher *β*) from our baseline parameterization (red star), the adaptive protocol with toxicity modulation is the only protocol that appears as optimal (protocol 4 in Figure 4A). As we move further to the right, we interestingly find that it is the daily protocol with toxicity modulation that can also extend TTP beyond the simulation period (protocol 3 in Figure 4A). Finally, increasing *β* further, moving to the extreme end of parameter space for which sensitive cells have a competitive advantage over resistant cells, the adaptive protocol joins the two toxicity protocols as all three prevent progression within the simulation period.

### 3.4 Rates of Drug-induced Death and of Toxicity Recovery Affect the Nature of Treatment Failure

It is quite natural to be interested in the *cause* of treatment failure. Surprisingly, simulations revealed that while *α*_*R*_ strongly influences time to failure, it does not play an important role in determining the cause of failure (data not shown). For this reason, we explored the underlying cause of progression by performing a parameter sweep in (*β*, *δ*)-space, where *δ* is the drug-induced death rate. For each protocol and each (*β*, *δ*) pair, keeping all other parameters fixed at their baseline value in Table 1, we compute the tumor composition (fraction of resistant cells) at the time of treatment failure. The results, shown in Figure 5, indicate that despite the importance of *β* in determining TTP, the role of *β* in the cause of treatment failure is minimal and can only be noticed in the protocols with toxicity at intermediate values of *δ*. In fact, the same can be said of *δ* in the protocols without toxicity, as Figure 5A,B shows that treatment failure consistently results from the eventual dominance of resistant cells, with sensitive cells nearly eradicated. However, as shown in Figure 5C,D, introducing toxicity feedback creates a region in parameter space where progression occurs due to the uncontrolled growth of sensitive cells. This occurs at low *δ* values, where drug efficacy is insufficient to control the sensitive tumor bulk.

**Figure 5.**
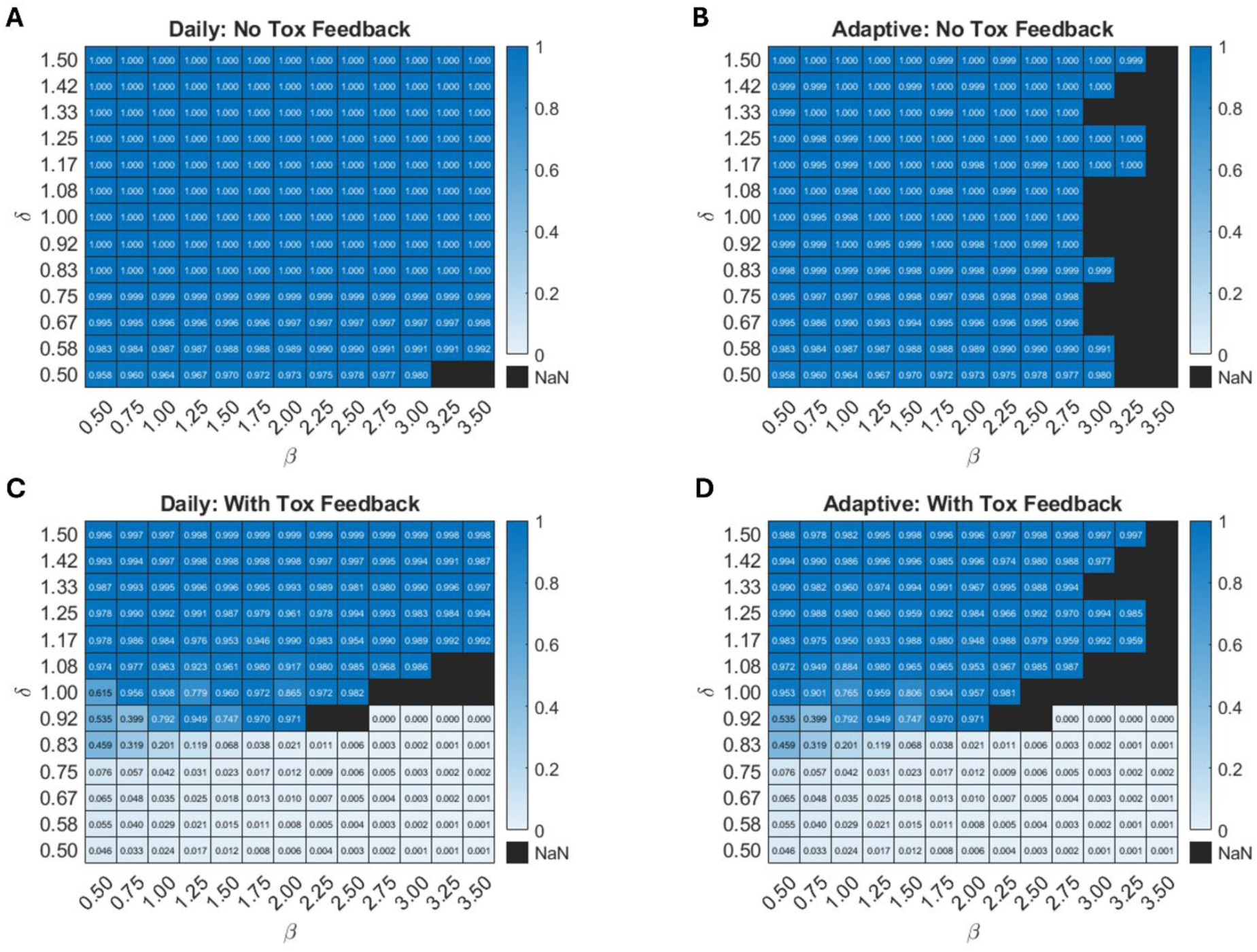
Fraction of resistant cells at TTP for different combinations of *β* and *δ* tested over the four protocols: (A) daily, (B) adaptive, (C) daily with toxicity, and (D) adaptive with toxicity feedback. All other parameters are fixed at their baseline values reported in Table 1. Black regions indicate that treatment did not fail within the simulation time of 100 days.

The global sensitivity analysis in Figure 2 identified the toxicity recovery rate *γ* as either the least sensitive, or second least sensitive, parameter in all four protocols. This is surprising, as one would think that the rate of toxicity recovery would affect the outcomes of protocols with toxicity feedback. If the rate of toxicity recovery does not significantly influence TTP, perhaps it affects the cause of treatment failure? We explore this question next.

Figure 6 assesses the effect of *γ* in two different parameter regimes identified from Figure 5C,D: when sensitive cells drive progression (*δ* = 2/3) and when resistant cells drive progression (*δ* = 4/3). In both parameter regimes, we observe a bifurcation in the reason for treatment failure. When *γ* is low, toxicity accumulates, forcing treatment breaks that allow sensitive cells to regrow and outcompete resistant cells, ultimately driving treatment failure. Conversely, when *γ* is high, toxicity resolves quickly, permitting more frequent treatment cycles that, in turn, eliminate sensitive cells leaving the resistant cells to drive treatment failure. The bifurcation in behavior happens at the same value of *γ* for each value of *δ* considered (compare Figures 6A and 6B with *δ* = 2/3, then compare Figures 6C and 6D with *δ* = 4/3).

**Figure 6.**
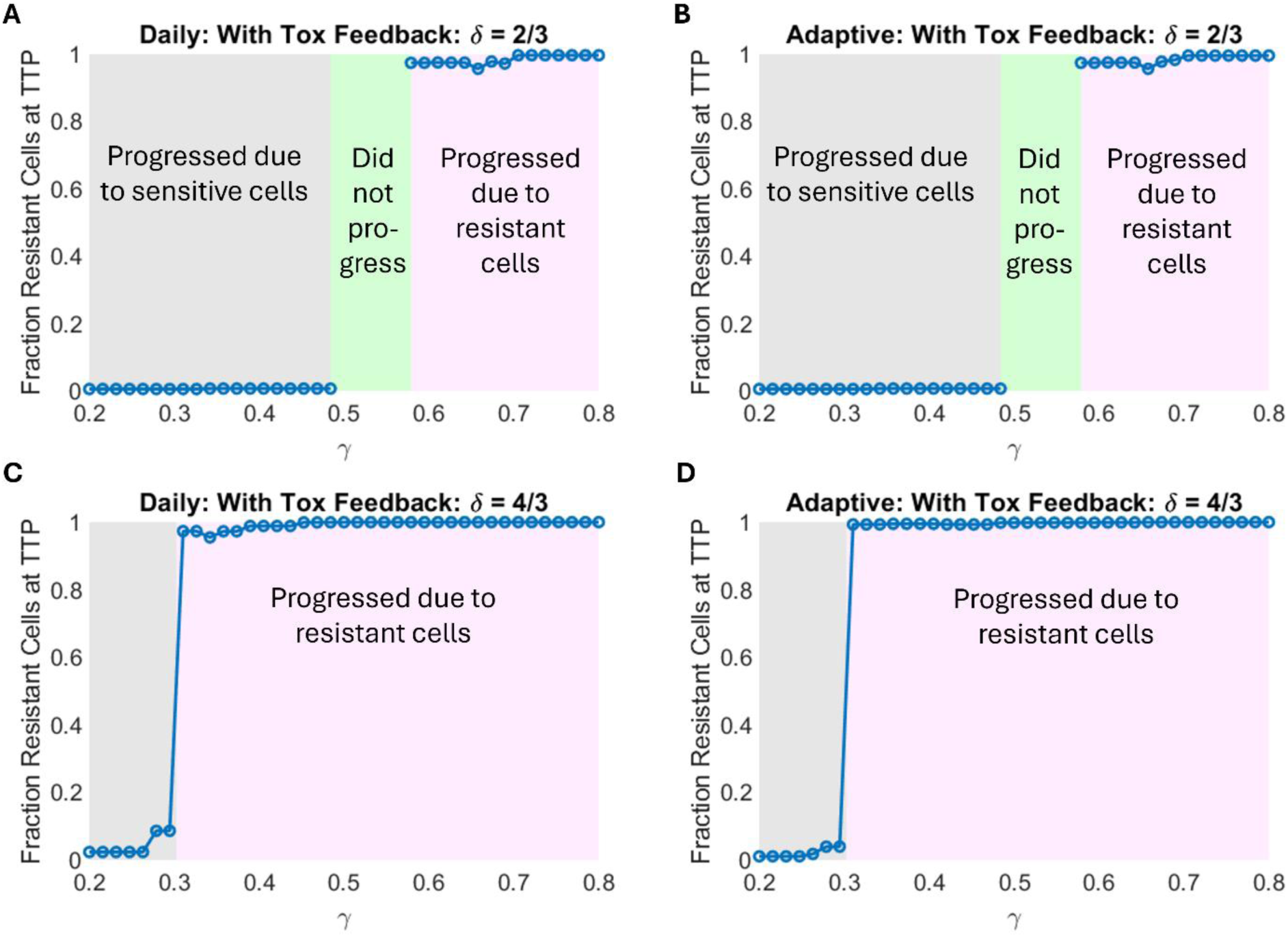
Effect of toxicity clearance, *γ*, on the nature of treatment failure. As *γ* increases, the reason for treatment failure switches from sensitive to resistant cells. (A) Daily with toxicity and (B) adaptive with toxicity protocols when drug effectiveness, *δ*, is small (*δ* = 2/3). (C) Daily with toxicity and (D) adaptive with toxicity protocols when *δ* is large (*δ* = 4/3). The remaining parameters are fixed at their baseline value in Table 1.

Interestingly, there is a range of intermediate *γ* values in the *δ* = 2/3 case where treatment does not fail within the simulation time, and in some cases, may result in prolonged cycling of therapy. In other words, there is a range of values for *γ* where toxicity is well controlled and with this moderate drug effectiveness, the feedback-modified treatments work well. However, at higher values of treatment effectiveness, *δ*, this ‘Goldilocks’ window vanishes, and treatment invariably fails during the simulation period. Prolonged cycling of treatment is no longer a possibility as resistant cells will lead to treatment failure. These results demonstrate that even though *γ* has a minimal effect on TTP (sensitivity analysis in Figure 3), it can play a crucial role in determining the cause of treatment failure.

### 3.5 Assessing Therapy Outcome Robustness to Protocol Thresholds

Next, we assess whether treatment outcomes can be improved by adjusting the therapy on/off thresholds in adaptive and toxicity-tracking protocols. We previously showed that at baseline tumor size and toxicity thresholds for starting or stopping therapy, TTP varies significantly with *β* and *α*_*R*_. Here, we fix all model parameters to the baseline values from Table 1 and instead vary the tumor size thresholds Rx_on_ and Rx_off_ in the adaptive protocols, or the toxicity Tox_on_ and Tox_off_ in the protocols with toxicity feedback. We report predicted TTP for each of the following cases: the effect of varying Tox_on_ and Tox_off_ in daily and adaptive protocols with toxicity feedback (Figures 7A and 7B, respectively) and the effect of varying Rx_on_ and Rx_off_ in adaptive protocols without or with toxicity feedback (Figures 7C and 7D, respectively).

**Figure 7.**
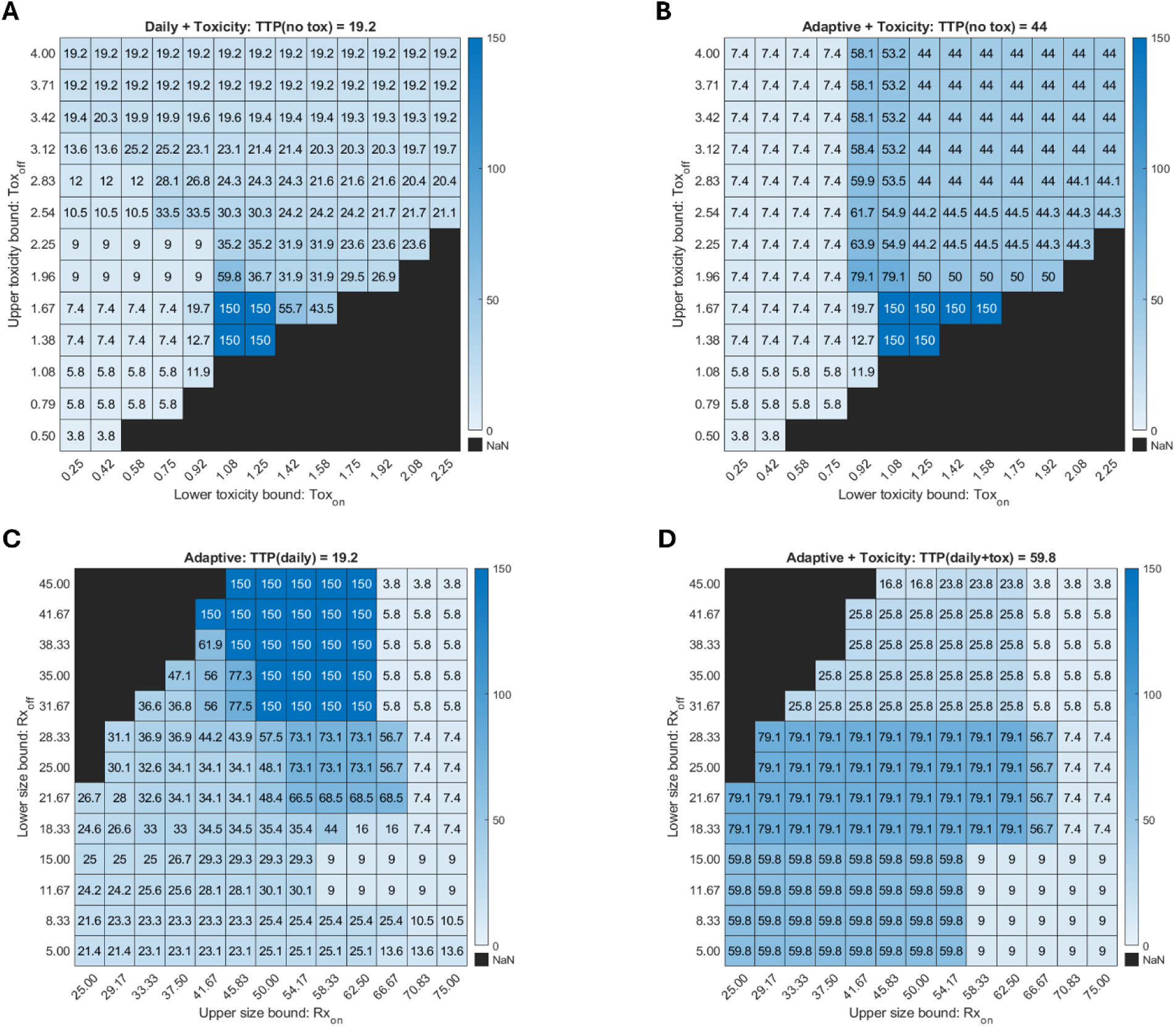
TTP as a function of protocol thresholds at baseline parameter values. Top row: toxicity thresholds varied in the daily with toxicity (A) and adaptive with toxicity (B) protocols. Bottom row: tumor size thresholds varied in the adaptive (C) and adaptive with toxicity (D) protocols. The results of the same protocol sweep for smaller and larger values of *β*, than the baseline value of *β* = 2.4 used here, are shown in Figure S3.

Figures 7A and 7B show TTP as a function of Tox_on_ (the toxicity level at which treatment may resume, horizontal axis) and Tox_off_ (the toxicity level at which therapy is paused, vertical axis). The constraint Tox_on_ < Tox_off_ must be satisfied; protocols violating this condition are blacked out in the bottom right corners. When Tox_off_ is too high, that is, when we are allowing a high level of toxicity in the simulation, toxicity-induced treatment pauses are infrequent. As a result, TTP in toxicity-feedback protocols is similar to that of protocols without toxicity feedback in both daily (TTP of 19.2 days) and adaptive (TTP of 44 days) dosing. Low values of Tox_on_, meaning treatment can only resume after toxicity has mostly resolved, result in sharp declines in TTP. In these cases, if toxicity triggers a treatment pause (as seen with moderate Tox_off_ values in the daily protocol, and all values in the adaptive protocol), the toxicity level never falls low enough for the treatment to restart. The result is that cancer cells grow unchecked and TTP is very short. Conversely, high values of Tox_on_ allow for treatment to be resumed sooner, minimizing the effect of toxic feedback on the protocols, and resulting in similar TTPs to that of protocols without toxicity feedback.

Notably, the longest TTP for daily and adaptive protocols with toxicity feedback occurs at intermediate toxicity thresholds that are fairly close together, forcing toxicity to remain relatively constant and moderate. Within this sweet spot in protocol space, the model predicts that the tumor will not progress during the 100-day simulation window. These protocols with an optimal TTP occur at the boundary of the blacked-out region (Figures 7A and 7B). Interestingly, this tumor control behavior occurs over a larger range of protocol thresholds for the adaptive with toxicity protocol, as compared to the daily with toxicity protocol.

Figures 7C and 7D show TTP as a function of Rx_on_ (the tumor size at which treatment resumes, horizontal-axis) and Rx_off_ (the tumor size at which treatment pauses, vertical-axis). The constraint Rx_on_ > Rx_off_ must be satisfied; parameter sets violating this condition are blacked out in the top left corners. Looking at the rightmost columns in Figures 7C and 7D, when Rx_on_ is too high, the tumor is allowed to grow to a sufficiently large size before a new treatment cycle begins. Under both adaptive protocols, TTP is noticeably lower than in the corresponding daily protocols (reported in the title of each subfigure), regardless of Rx_off_. When Rx_on_ is small, both adaptive protocols (with or without toxicity feedback) are comparable to, or modestly outperform, the corresponding daily protocol.

As we saw when varying the toxicity thresholds, there is a sweet spot in the adaptive protocol space for which the tumor is controlled over the 100-day window. This optimal window is found at high Rx_off_ and intermediate Rx_on_ values (indicated by the boxes labeled ‘150’ in Figure 7C). This region suggests that the best outcome is obtained by pausing treatment before the tumor shrinks too much, thereby allowing the sensitive cell population to recover and suppress resistant cells through competition, and by restarting treatment before the tumor becomes too large.

Interestingly, we do not observe the same sweet spot in protocol space for the adaptive therapy with toxicity feedback (Figure 7D). At high Rx_off_ and intermediate Rx_on_ values, TTP is smaller than what we observe with the baseline threshold values (59.8 days). Instead, an optimal treatment window emerges at intermediate Rx_off_ values, where tumor burden is maintained between 36.6% and 56.6% of its initial size. Although adjusting size-based thresholds can improve TTP, none of the tested protocols produce a durable response – defined here as TTP exceeding the 100-day simulation period – once toxicity feedback is included in the adaptive protocol. While we did not identify tumor size thresholds for which the adaptive protocol with toxicity can control the tumor in Figure 7D, Figure 7B shows that toxicity feedback can produce the desired outcome. Both Figures 7B and 7D are looking at two-dimensional slices of a four-dimensional protocol space, so it is important not to interpret either figure along the lines that the adaptive protocol with toxicity feedback is inferior to the others, as these are snapshots of a larger space.

The threshold sweeps in Figure 7 also highlight the personalization level required to optimize the toxicity feedback protocol. We see that, depending on the chosen toxicity thresholds, the efficacy of a toxicity feedback protocol can improve or worsen relative to the daily protocol’s TTP of 19.2 days. As an example, consider Figure 7A and fix the maximum tolerated toxicity at Tox_off_ = 1.67. Moving the treatment restart threshold Tox_on_ from 0.92 to 1.08 results in a jump in TTP from 12.7 days (worse than the daily protocol) to over 100 days (much better than the daily protocol). However, a further increase of Tox_on_ reduces the predicted TTP, revealing the existence of a toxicity threshold “sweet spot” for this fixed parametrization. Such “sweet spots” exist for all the treatment cycling protocols, and the landscapes will depend on the model parameters.

### 3.6 Optimizing Threshold “Sweet Spots” Across a Virtual Population

Finally, to systematically determine whether adaptive and toxicity-based thresholds can be optimized for maximal TTP across various parameter values, we conduct a virtual patient analysis. In this approach, each “virtual patient” is assigned a unique set of values for five parameters, *α*_*S*_, *ϵ*, *β*, *δ*, and *γ*, drawn from independent lognormal distributions (Figure S4). The means of these distributions match the corresponding baseline values in Table 1, and the standard deviations are chosen to approximate the parameter ranges used in the sensitivity analysis of Figure 3. We then repeat the protocol threshold analysis from Figure 7 for each virtual patient and calculate the mean TTP across the entire virtual population, for every pair of threshold values. As before, VPs that do not reach treatment failure in 100 days are assigned a TTP of 150. The results for 100 simulated VPs are reported in Figure 8.

**Figure 8.**
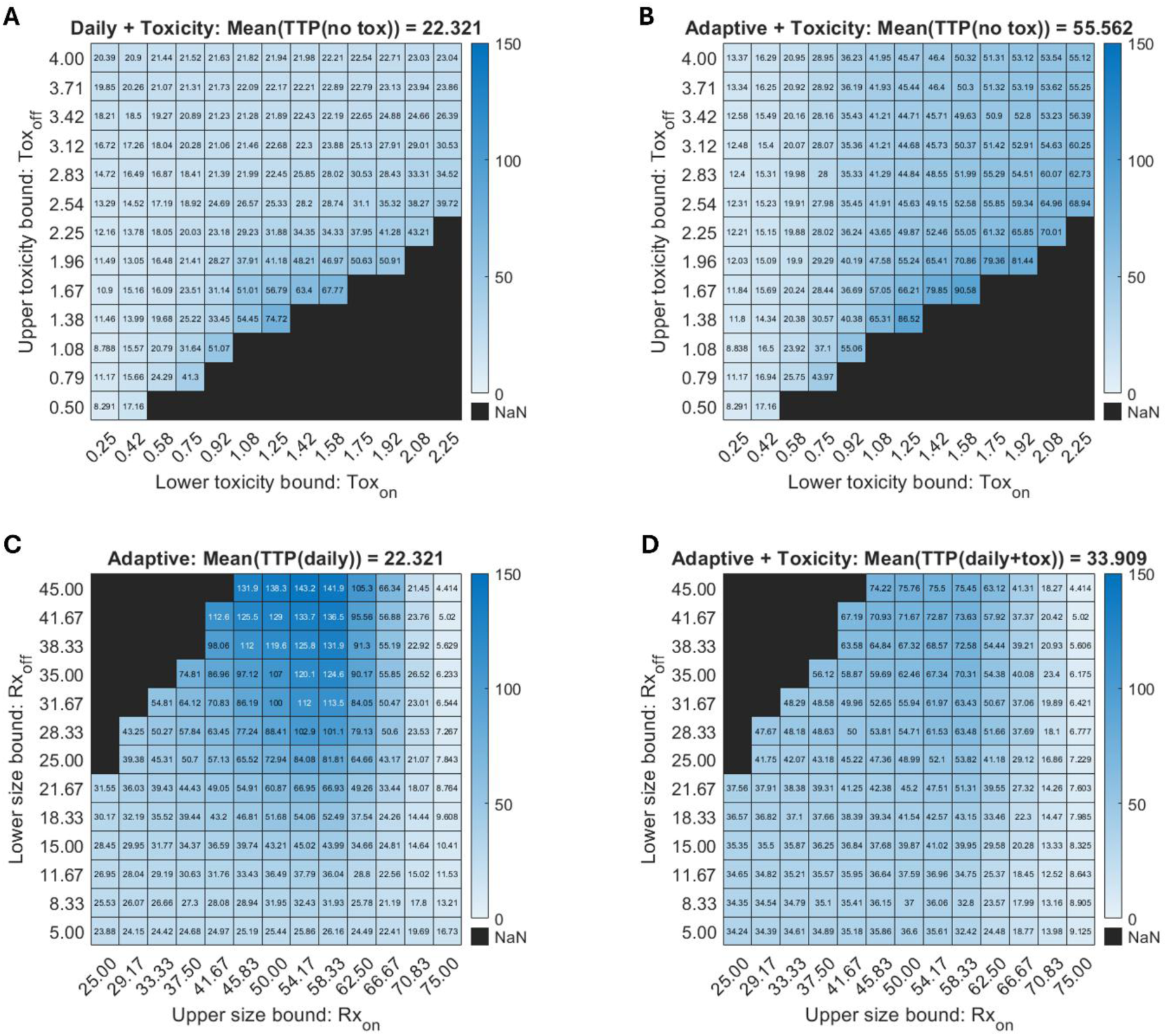
Mean TTP for 100 VPs. Top row: toxicity thresholds varied in the daily with toxicity (A) and adaptive with toxicity (B) protocols. Bottom row: tumor size thresholds varied in the adaptive (C) and adaptive with toxicity (D) protocols.

Figures 8A and 8B show mean TTP as a function of Tox_on_ (the toxicity level at which treatment resumes, horizontal-axis) and Tox_off_ (the toxicity level at which treatment pauses, vertical-axis), with the tested protocol specified in the panel title, along with the corresponding mean TTP for the protocol without toxicity feedback. Again, the blacked out bottom right corners correspond to pairs of values that violate the constraint that Tox_on_ < Tox_off_. Figures 8C and 8D show mean TTP as a function of Rx_on_ (the tumor size at which treatment resumes, horizontal-axis) and Rx_off_ (the tumor size at which treatment pauses, vertical-axis), with the tested protocol specified in the panel title, along with the corresponding mean TTP for the protocol without tumor size-based feedback. The upper left corner is blacked out as this region violates the constraint that Rx_on_ > Rx_off_.

It is instructive to compare the heatmaps in Figure 7 for the baseline parameterization to the heatmaps in Figure 8, where we are looking at average behavior across a virtual population. When the toxicity thresholds are varied, we generally see that the sweet spot in protocol space is comparable between the baseline parameterization (Figures 7A and 7B) and the virtual population (Figures 8A and 8B). The same is true when the tumor size thresholds are varied in the adaptive protocol (Figure 7C versus Figure 8C). Interestingly, the analogous consistency is not observed in the adaptive protocol with toxic feedback (Figure 7D versus Figure 8D). At the baseline parameter values, intermediate Rx_on_ values coupled with large Rx_off_ values corresponded to a suboptimal treatment protocol. However, this same region of protocol space is optimal for the average of the virtual population. This demonstrates that the effect of tumor size thresholds in the adaptive with toxicity protocol depends, to a large extent, on the model parameters.

To demonstrate the variability across VPs, as opposed to the average behavior as in Figure 8, we examine the distribution of patient outcomes by plotting Kaplan-Meier curves (Figure 9). These curves illustrate the proportion of virtual patients who have not progressed over time. The Kaplan-Meier curve for each protocol uses the thresholds that were identified to optimize the mean TTP for the virtual population in Figure 8. For instance, the adaptive protocol in Figure 8C has the longest TTP when Rx_on_ = 58.33 and Rx_off_ = 45, so these are the thresholds set for the adaptive protocol in Figure 9. For reference, we also include the Kaplan-Meier curve for the daily protocol with no treatment pausing (blue curve). The adaptive with toxicity protocol (purple curve) in Figure 9 uses the optimal thresholds found by searching over the four-dimensional space defined by the two toxicity thresholds and the two tumor size thresholds.

**Figure 9.**
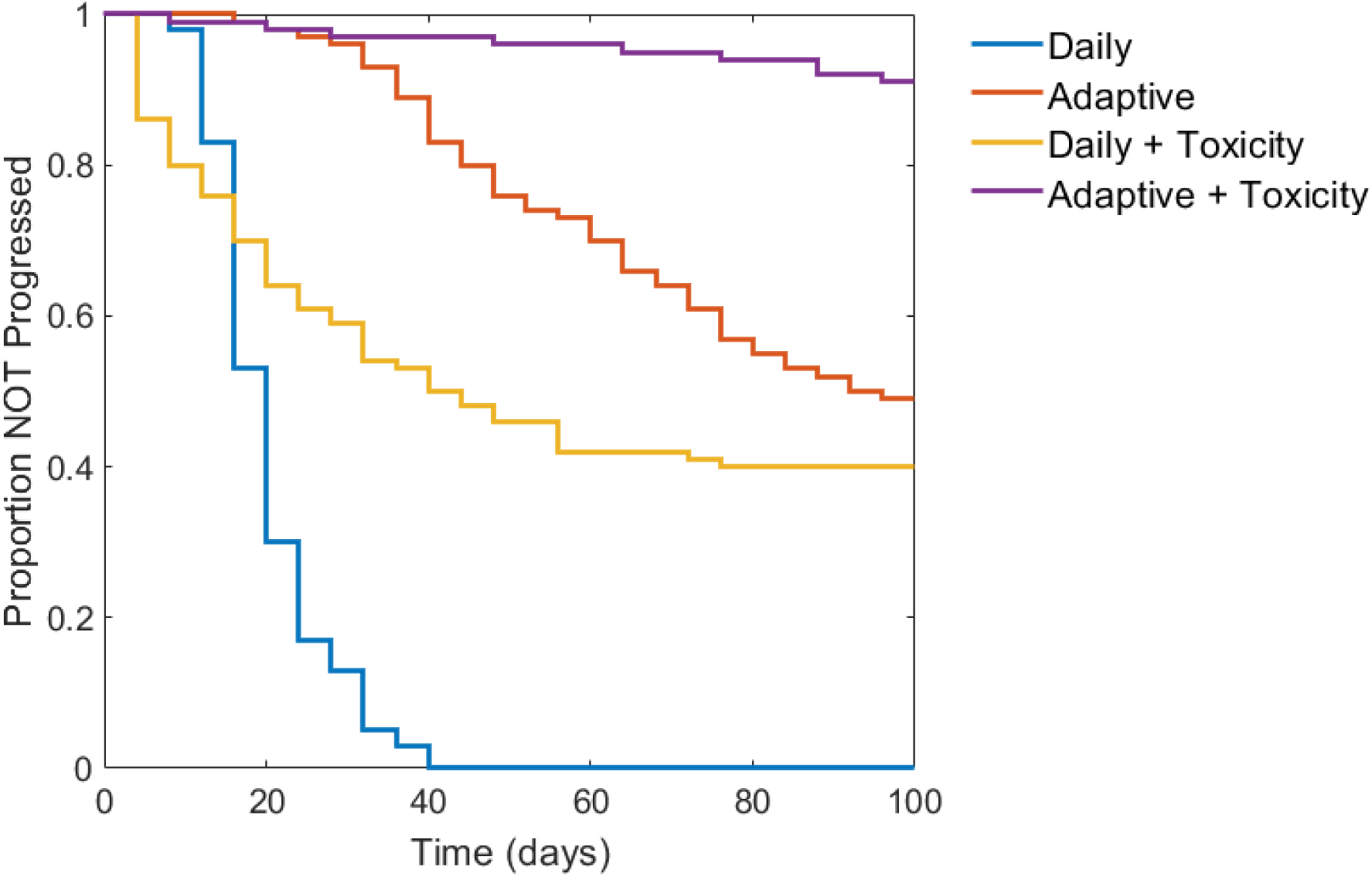
Variability across a virtual population for each protocol with optimized treatment thresholds. Mean TTP of 100 VPs was used to determine the optimal thresholds for each protocol (except the daily protocol which does not have any thresholds).

By adding toxicity considerations to the daily protocol (yellow curve in Figure 9) 40% of VPs experience a TTP greater than 100 days, but 16% experience a TTP shorter than the daily protocol. The adaptive protocol (red curve) significantly improves TTP for the majority of VPs compared to the daily protocol and leaves almost 50% of VPs with no disease progression by 100 days. Finally, the adaptive with toxicity protocol (purple curve), demonstrates the potential advantage of dual feedback: 91% experience no progression by 100 days.

## 4 Discussion

Cancer treatments can fail for many reasons that are not necessarily the loss of treatment efficacy or the emergence of resistance. For instance, sometimes treatments fail due to unacceptable levels of toxicity. Previous mathematical models suggest that adaptive therapy (i.e., pausing and restarting treatment based on tumor size) can improve patient outcomes by delaying the emergence of resistance. However, existing methods that rely on tumor burden as the sole feedback mechanism fail to capture the significant impact of treatment-related toxicity, a critical factor in clinical practice. Toxicity often necessitates dose reductions or treatment breaks. Current adaptive therapy clinical trials use toxicity management guidelines, often adapted from MTD trials, but do not explicitly incorporate toxicity thresholds into the treatment design. The aim of this study was to investigate how the addition of toxicity constraints into a model of adaptive therapy may influence TTP.

We demonstrated that toxicity constraints can enhance or diminish outcomes depending on parameter values and protocol thresholds. In our analysis, we compared TTP of four protocols: daily; adaptive, where treatment is controlled based on tumor size criteria; daily with toxicity feedback, where treatment is controlled based on accumulated toxicity criteria; and adaptive with toxicity feedback, where treatment is controlled by both tumor size and toxicity criteria. We were able to identify scenarios in which each of the four protocols could optimize TTP. The optimal protocol depended on the growth rates of, and competition between, the sensitive and resistant cells, and on the defined protocol thresholds. Protocols leading to the longest TTPs typically incorporate either or both efficacy and toxicity feedback rules.

Additionally, by assessing tumor composition at TTP, we found that there are two paths to treatment failure. The tumor may progress due to significant growth of either sensitive or resistant cells, with progression due to sensitive cells occurring only in protocols with toxicity feedback. As such, drug dose timing becomes critically important: waiting too long to resume treatment (for example, because of slow toxicity recovery), may result in sensitive cell growth causing treatment failure.

Our results suggest that varying toxicity and efficacy thresholds can significantly affect treatment response, with relatively small changes in threshold values potentially having significant effects on TTP. We also observed that TTP is sensitive to several model parameters. For these reasons, we conducted a virtual population analysis to better understand the relationship between protocol thresholds and TTP across a heterogeneous population. In particular, we varied the protocol thresholds for both efficacy and/or toxicity (as relevant to the protocol), to determine the optimal thresholds leading to the longest average TTP for the virtual population. We found that, while there do not exist thresholds that guarantee lack of progression for the entire population, there do exist thresholds that can significantly extend TTP (on average) for all protocols with feedback. The longest average TTP, with the most non-progressing VPs, was predicted for the dual-feedback protocol incorporating both adaptive and toxicity constraints, as demonstrated by the simulated Kaplan-Meier survival curves in Figure 9.

While these predictions are based on a conceptual model, emerging clinical data does lend support to the notion that treatment modulation based on toxicity constraints may not be inferior in efficacy to more traditional approaches. For instance, in (36), the authors report that patients with metastatic renal cell carcinoma who received fewer than four doses of ipilimumab and nivolumab due to reasons other than disease progression (primarily toxicity) had similar overall survival rates at 18 months to those who completed all four doses. In fact, the median overall survival rate was actually larger (82.5 months versus 67.1 months) in the patients who received fewer than four doses (36).

These results align well with the principles of adaptive therapy, where treatment pauses based on tumor size (though not toxicity) can actually maintain or even improve efficacy (37), potentially through mitigating competition between sensitive and resistant cancer cells. By pausing or reducing treatment upon reaching specific toxicity thresholds and resuming once resolved, patients may experience fewer adverse effects without compromising treatment outcomes. The importance of such an approach is further supported by the FDA’s Project Optimus (38), where drug developers are encouraged to move away from a “no regrets” MTD-like strategy towards a more nuanced, albeit harder to develop, protocol that balances efficacy and toxicity.

Psychological resistance, both among patients and clinicians, may pose a hindrance in the potential clinical implementation of pausing therapy due to toxicity, as it may be seen to imply a lack of commitment or a weaker fight against cancer. This stigma may arise from multiple factors, such as the implicit belief that more aggressive treatment leads to a better outcome (the “more is better” mindset) and patient expectations and fears, where patients associate pausing treatment with disease progression and failure. It is also possible that the image of a cancer patient as a “fighter”, while encouraging, does not leave room for managing the immense physical and psychological challenges of cancer treatment without sacrificing the image. Communication of the benefits of a potential toxicity-monitoring protocol would thus be necessary. It should be emphasized, when discussing the treatment protocol, that pausing therapy is intentional, designed to maximize long-term benefit, and not a sign of weakness.

It should be noted that the model used here is conceptual in nature: it uses a simplified description of tumor heterogeneity, a simplified representation of toxicity as a function of drug concentration only, and it excludes many tumor microenvironmental and immune factors. Furthermore, it uses a one-compartment pharmacokinetic model, ignoring any variability in drug kinetics or dose amounts. Finally, toxicity thresholds values were chosen to illustrate differences between the four compared protocols.

In future work, some of these aspects will be expanded upon, with pharmacokinetic properties of specific drugs and a more refined approach to quantifying toxicity and modulating dose in response to it. For instance, a frequent side effect of many cytotoxic drugs is neutropenia, which is associated with increased risk of infection. Neutropenia is measured by absolute neutrophil count. The pharmacokinetics of a specific drug can be coupled with neutrophil quantification, improving the direct translatability of the model. That said, patients do experience a spectrum of side effects (fatigue, nausea, weakness, muscle loss, blood count drops, etc.), and deciding how to aggregate these into decision rules will require careful consideration. Adequately quantifying a composite toxicity score in real time is a complex task and may require multiple criteria-based constraints.

In conclusion, our modeling study demonstrates how incorporating toxicity constraints into adaptive therapy can reduce treatment-associated toxicity without sacrificing efficacy. By balancing patient tolerability with treatment efficacy, such toxicity-informed adaptive protocols hold the promise of turning at least some types of cancer into manageable chronic conditions. This approach may extend survival rates, improve patient quality of life, and potentially bring us closer to truly personalized cancer treatment.

## Acknowledgements

This work was initiated during the Thematic Program on Mathematical Oncology at the Fields Institute for Research in Mathematical Sciences in Toronto, Canada. The authors thank Dr. Helen Byrne for insightful discussions during the workshop.

## Data Availability

Data sharing is not applicable to this article as no datasets were generated directly during the current study. Programming scripts in MATLAB are freely available at https://github.com/jgevertz/toxicity.

## Author Contributions

MD proposed the research question. All authors were involved with the initial project formulation. JLG, HVJ, IK and KPW developed the protocol algorithms, wrote the code, ran numerical simulations, designed figures, and prepared and edited all manuscript drafts. All authors contributed to editing and reviewing the manuscript.

## Supplementary Information

**Figure S1.**
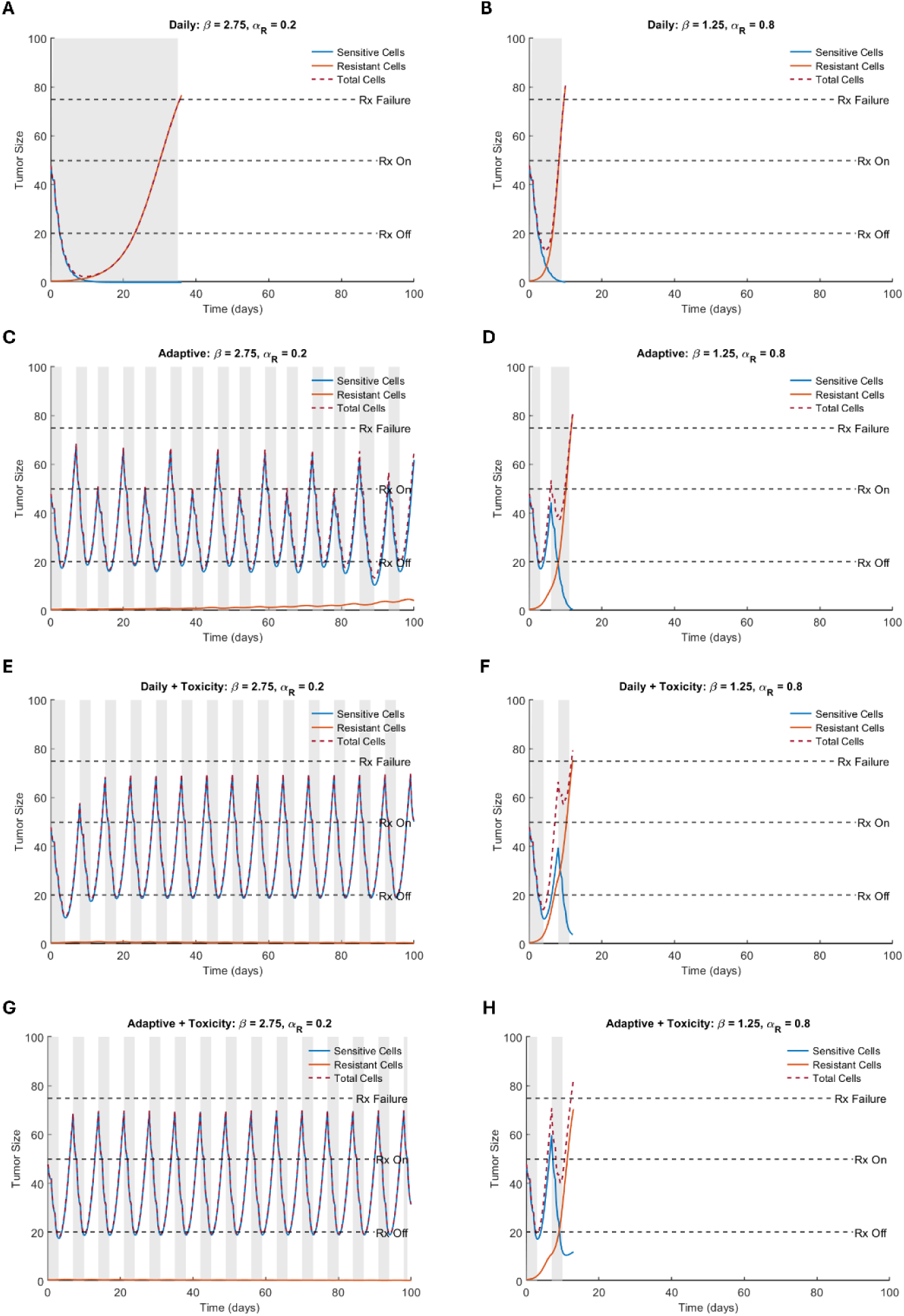
The four treatment protocols and resulting model outputs. The left column shows a parameterization for which resistant cells are slow growing but have a strong competitive advantage. The right column shows a parameterization for which resistant cells are fast growing but have a weaker competitive advantage. Compared protocols are daily (A-B), adaptive (C-D), daily with toxicity feedback (E-F), and adaptive with toxicity feedback (G-H). Time periods shaded grey represent when the treatment is on.

**Figure S2.**
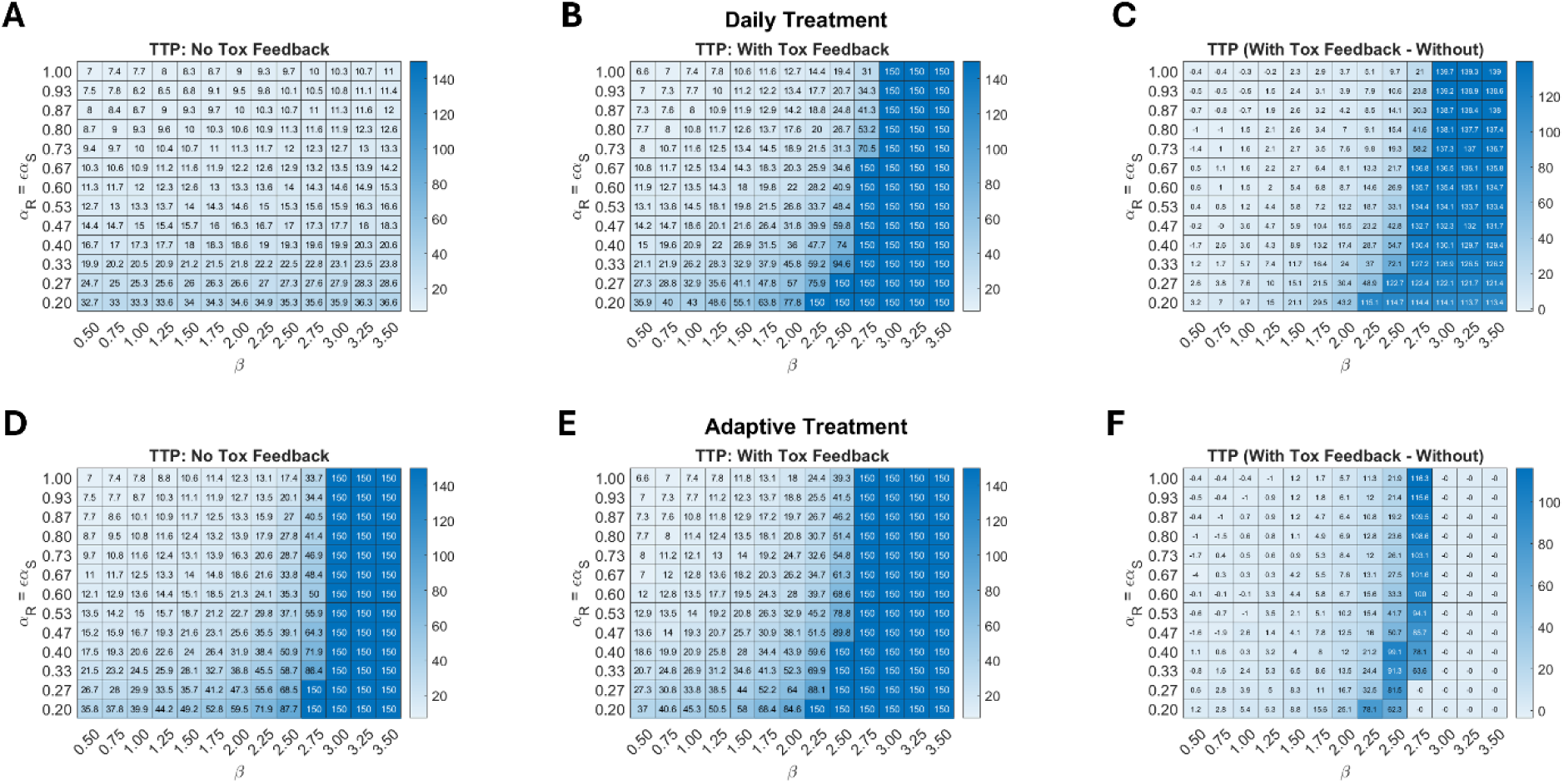
Parameter sweeps over *β* − *α*_*R*_ space. TTP is shown across parameter space for (A) daily protocol, (B) daily protocol with toxicity feedback, (D) adaptive protocol, (E) adaptive protocol with toxicity feedback. A value of 150 indicates that the predicted tumor did not progress within the simulation time of 100 days. The remaining parameters are fixed at their baseline value in Table 1. The right column shows the difference in TTP comparing the protocol with toxicity feedback to the same protocol without toxicity feedback: (C) daily, (F) adaptive.

**Figure S3.**
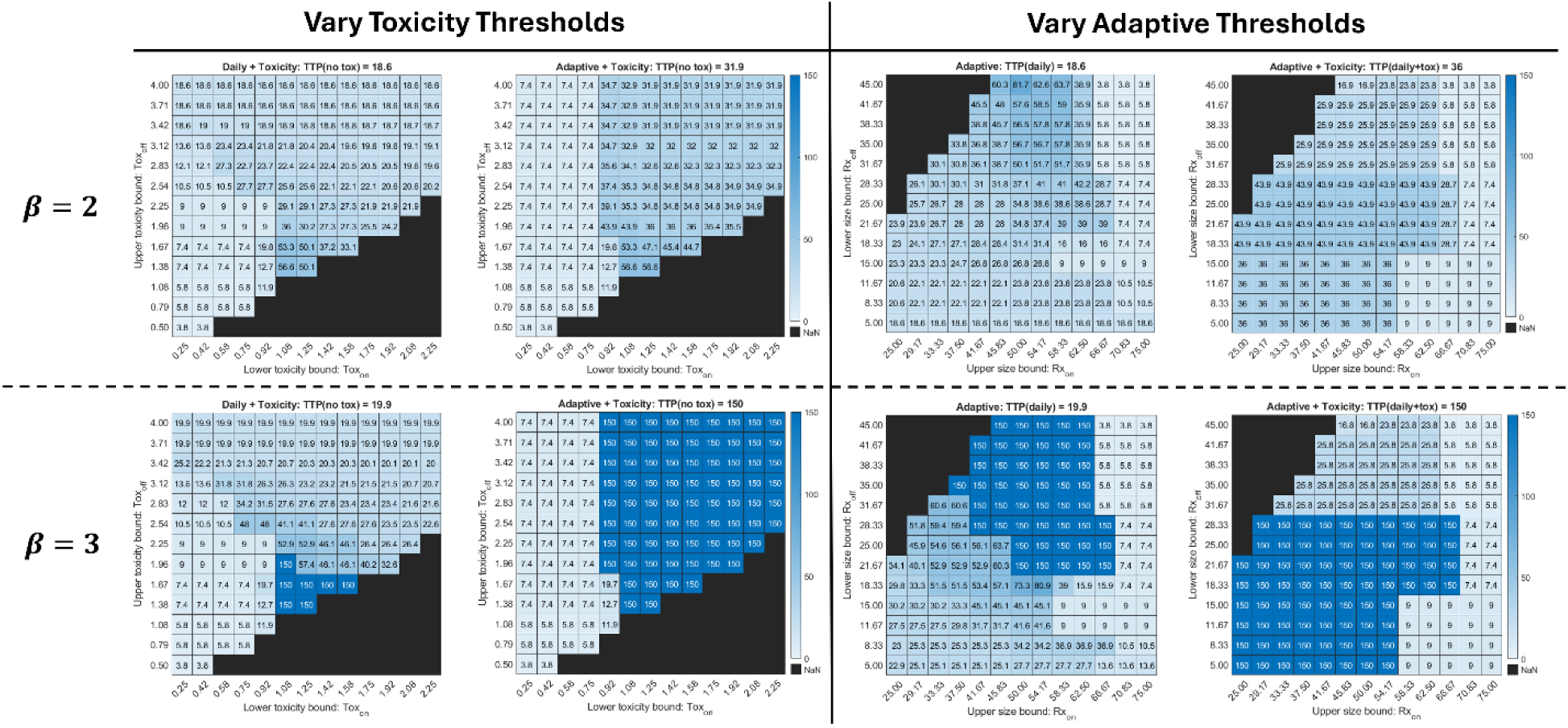
Effect of *β* on protocol sweeps.

**Figure S4.**
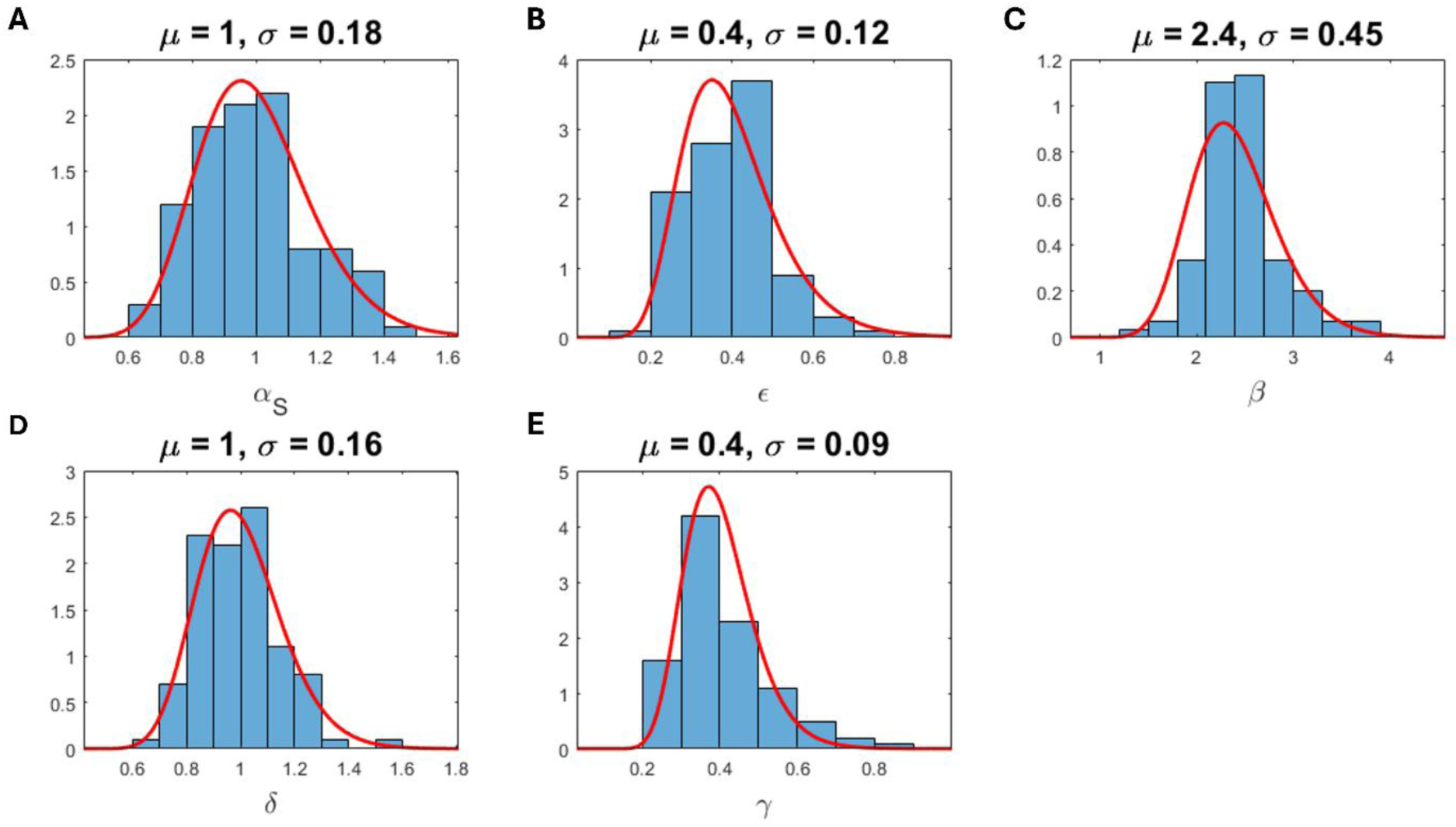
Lognormal distributions defined for each parameter of the VP analysis. The parameters defining the lognormal distribution (*μ* and *σ*) for each VP parameter (*α*_*S*_, *ϵ*, *β*, *δ*, and *γ*) are chosen such that the peak aligns with the baseline value reported in Table 1, and the width covers the range used in the global sensitivity analysis of Figure 3.

